# Deep attention based variational autoencoder for antimicrobial peptide discovery

**DOI:** 10.1101/2022.07.08.499340

**Authors:** Mahdi Ghorbani, Samarjeet Prasad, Bernard R. Brooks, Jeffery B. Klauda

## Abstract

Antimicrobial peptides (AMPs) have been proposed as a potential solution against multiresistant pathogens. Designing novel AMPs requires exploration of a vast chemical space which makes it a challenging problem. Recently natural language processing and generative deep learning have shown great promise in exploring the vast chemical space and generating new chemicals with desired properties. In this study we leverage a variational attention mechanism in the generative variational autoencoder where attention vector is also modeled as a latent vector. Variational attention helps with the diversity and quality of the generated AMPs. The generated AMPs from this model are novel, have high statistical fidelity and have similar physicochemical properties such as charge, hydrophobicity and hydrophobic moment to the real to the real antimicrobial peptides.

## Introduction

Antimicrobial resistance causes ∼2.8 million resistant infections yearly which leads to more than 700,000 deaths globally. This is expected to rise to 10 million deaths per year by 2050 if the current trend continues.^1–3^ Of particular importance is Multi-drug resistant Gram-negative bacteria. Naturally occurring antimicrobial peptides (AMPs) have remained effective to combat pathogens despite their ancient origins and continuous contact with pathogens. Therefore, AMPs are deemed as “drugs of last resort” for their ability to combat multi-drug resitant bacteria. AMPs are usually 12-50 amino acids long and are typically rich in cationic residues (R and K) as well as hydrophobic (A, C and L) amino acids. The mechanism of action of AMPs depends on their sequence but they generally act by disrupting the membrane or through other routes such as binding to DNA and essential cytoplasmic protein and inhibiting their function.^4^

There have been numerous studies on generating new AMPs and/or improving their activity which resulted in some successful AMPs.^5,6^ These have been generated usually through expert knowledge and rational design approaches which could be very costly due to vast space of peptides. There are some limitations in using current AMPs such as their relatively low half-lives, unknown toxicity to human cells, and relatively high production costs.^7–9^ On the other hand, due to the vast space of peptide sequence, computational techniques are necessary for discovery of novel AMPs with desired properties. Generative models in artificial intelligence have previously shown great promise in material and drug discovery.^10–12^ Deep learning have been previously used in peptide identification, property prediction and peptide generation.^13^ Specifically, deep generative models have been used for generating antimicrobial, anticancer, immunogenic and signal peptides to name a few. Computational methods using recurrent neural networks (RNNs),^14^ VAEs^15^ and generative adversarial networks (GANs)^16,17^ showed the promise of these methods for AMP discovery in silico.

In this study, we use variational autoencoders (VAEs) to learn a meaningful latent space of AMPs and generate novel AMPs from this latent space. A variational autoencoder^18^ encodes data into a latent space and decodes it back to the original data and optimizes a variational lower bound of the log-likelihood of the data. Since we are dealing with sequences as done in natural language processing (NLP) we use recurrent neural networks (RNNs) as both encoders and decoders in what is known as sequence-to-sequence models. Due to complexity of natural language and sequential nature of data, these models are harder to train than other types of neural networks. However, Bowman et al.^19^ showed that a seq-to-seq VAE is able to generate meaningful and novel sentences from the learnt continuous latent space. Attention mechanism proposed for translation originally, has made a great leap in NLP tasks.^20^ In attention mechanism, source information is summarized into a context vector using a weighted sum, where the weight are learned probabilistic distribution. This context vector is used during the decoding process to guide the decoder into what word in the sequence was most important during decoding. Attention was shown to significantly improve almost every task in seq2seq models such as translation,^21^ summarization,^22^ etc. However, Bahuleyan et al.^23^ showed that using a deterministic attention where the source information is directly provided during decoding can lead to a phenomenon called the “bypassing” where the variational latent space is not meaningful since the attention mechanism is too powerful. Thus they proposed a variational attention mechanism to address this problem where the attention vector (context vector) is modeled as a random variable by imposing a prior Gaussian distribution. They evaluated this model on question generation and dialog systems and showed that the variational attention achieves a higher diversity than deterministic attention while retaining high quality of generated sentences. In this study, we have used a variational attention with variational autoencoder in a seq2seq approach to generate novel, high quality and diverse AMPs. Moreover, we trained a binary classifier network using attention mechanism for evaluation of the generated peptides in the generative model. The generated peptides are also analyzed for their physicochemical properties and comparison with real antimicrobial peptides.

## Methods

The training data for the AMP prediction model was set by combining AMPs from multiple databases. These include DRAMP,^24^ LAMP2,^25^ DBAASP^26^ and APD3.^27^ All AMP sequences had a length of 5-30 amino acids. To exclude repetitive sequences from our dataset we used CD-HIT^28^ with a cutoff of 0.35 which resulted in 16808 AMP sequences. Since there are no known dataset for non-AMPs we made a non-antimicrobial dataset using Uniprot^29^ excluding keywords antimicrobial, antibiotic, antibacterial, antiviral, antifungal, antimalarial, antiparasitic, anti-protist, anticancer, defense, defensin, cathelicidin, histatin, bacteriocin, microbicidal and fungicide. The final non-AMP dataset had 16808 examples. We also ensured the positive and negative dataset have similar length distribution to avoid bias. he final code for AMP prediction and generation can be found at the github.com/ghorbanimahdi73/AMPGen.

### AMP prediction model

We trained a model on both AMP and non-AMP dataset for antimicrobial prediction. The architecture of our model is shown in figure 1A. The Antimicrobial classification network contains an embedding layer, a 2D convolution and a bidirectional LSTM with a context attention and a sigmoid activation at the end for binary classification of peptide sequences. The dataset was split into training (70%) and validation set (30 %). The output of the prediction model is the probability score for sequences. The sequences with score > 0.5 are considered AMP and those with score <0.5 are considered non-AMP. We used a binary cross-entropy loss and an Adam optimizer for training the network. Early stopping was applied if the validation loss was not improving for 5 consecutive epochs during training. The weights of the model with the best validation accuracy was selected as the optimal model. A 10-fold cross-validation was applied to tune the hyperparameters of the model. In the AMP prediction network, the peptide sequences are first transformed into a sequence of integers from 1 to 20 which are then embedded into 2d matrices in the embedding layer of the network. The optimal embedding size was found to be 64 dimension. A 2D convolution is then applied to the embedded sequences using 64 convolutional filters of size 3. The output of the convolutional layer then goes into a bidirectional-LSTM which processes the matrices for each residue from both forward and backward directions and the output is the summation of the two directions. The tuned bi-LSTM hidden dimension was 64. The attention layer then gather the hidden state of bi-LSTM and computes a weighted sum of all hidden states as:

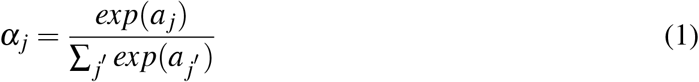

where *a* is the attention score by applying a linear transformation to the bi-LSTM outputs followed by a ReLU activation and *α* is the attention weight. The output of the attention layer can be computes a a weighted sum ∑ _*j*_*α*_*j*_*h*_*j*_ where *h*_*j*_ is the *j*’th hidden state of the bi-LSTM output. The output of attention then goes into the sigmoid activation for antimicrobial prediction. The final model had 93% accuracy under a 10-fold cross-validation which is comparable to other AMP-prediction models such as AMPlify and ACEP with 93.7 % and 92.6% accuracies. However, the goal of this study is not antimicrobial prediction and this network was trained in order to evaluate the generated peptides by our generative model.

**Figure 1:**
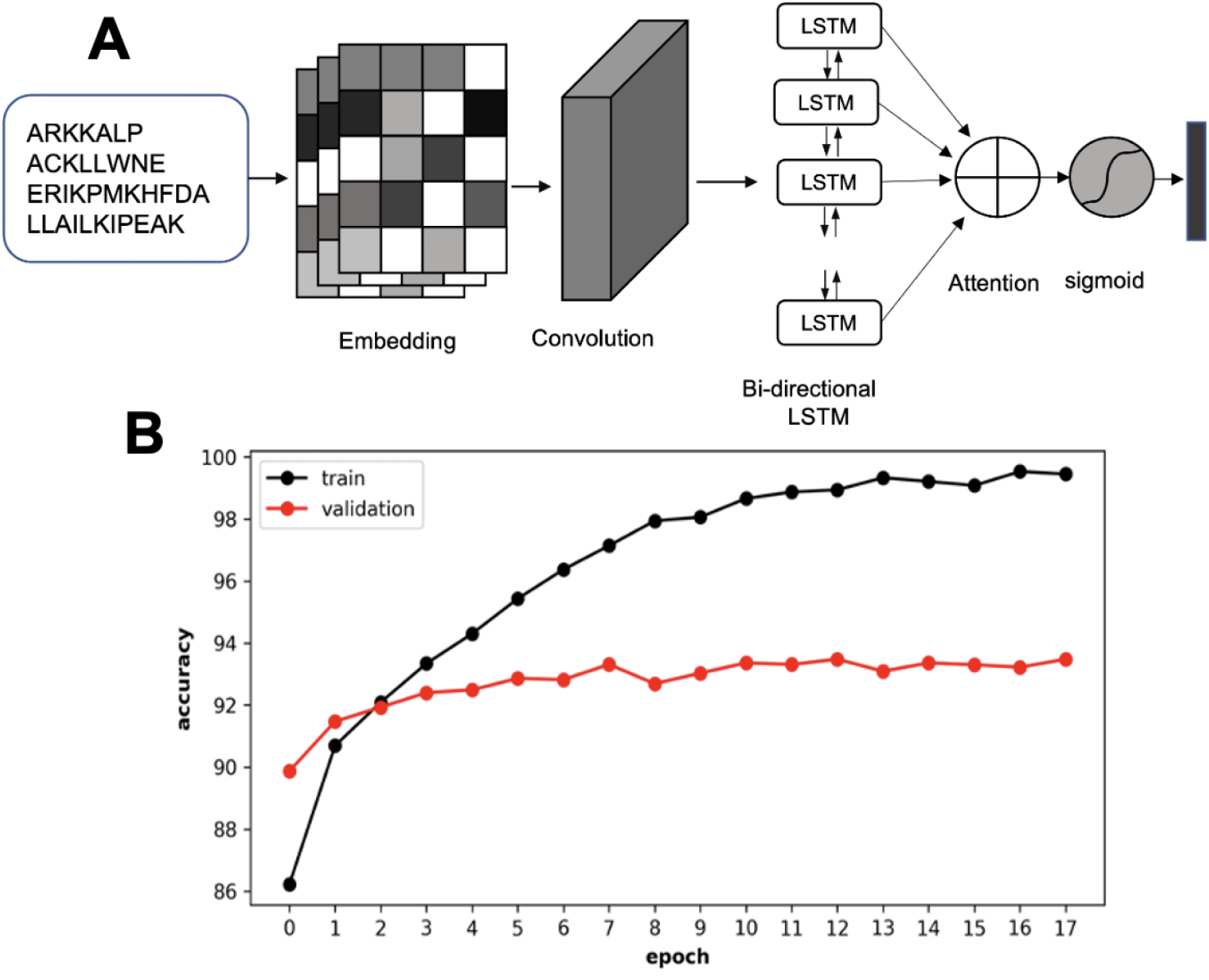
A) An illustration of the classification network used for evaluating the generated AMPs. B) training and validation accuracy of AMP prediction model

### Variational autoencoder

A traditional VAE proposed by Kingma and Welling encodes data to a latent space and then decodes to reconstruct the input data.^18^ The network is trained to optimize the variational lower bound of the log-likelihood of data. Since we are dealing with sequences, in natural language processing (NLP) recurrent neural networks (RNNs) are typically used as encoders and decoders in what is known as sequence to sequence models (seq2seq). Bowman et al^19^ trained a seq2seq VAE and used the continuous latent space to generate new text. A model with useful information in the latent space will have non-zero KL and a relatively small cross-entropy term. However, in a standard VAE, the KL becomes vanishingly small and the model becomes a RNN language model. The decoder learns to ignore the latent *z* vector and only use the input data which is provided at each step of decoding. Two techniques have been proposed by Bowman et al. to mitigate these issues, both of which are used here: 1) KL-annealing and 2) word dropout. For the KL-annealing, we add a variable weight to the KL term in the loss function during training. This weight is set near zero at the beginning of training and then increases to a maximum weight toward the end of training process. This ensures that the at the beginning the model learn enough information from latent space. The word dropout weakens the decoder by removing some of the conditional input information during training. This forces the model to rely more on the latent code.

Attention mechanism has transformed the natural language processing enabling training of enormous models and achieving high accuracies. In the attention mechanism, the source information is summarized in an attention vector using a weighted sum of hidden states of the source sentence where the weights are learned probabilistic distribution. Then this attention vector is directly fed to the decoder at each step during decoding. It has been demonstrated that this attention mechanism improves the performance of models in translation,^21^ Summarization^22^ and other NLP tasks. However, it was shown that this deterministic attention can serve as a bypassing mechanism and the latent space cannot learn the distribution of the data since the attention is too powerful.^23^ Here we use variational auto-encoder with variational attention for the generation of novel antimicrobial peptides. Different parts of the network are described in detail below:

### Encoder

In our model, encoder is a GRU which is parameterized by *θ*_*E*_. The encoder network takes the inputs sequence *x* = *x*_1_, …, *x*_*n*_ and outputs the hidden representation of the sequence *h* = *h*_1_, …, *h*_*n*_ where n is the length of the sequence. This is written as :

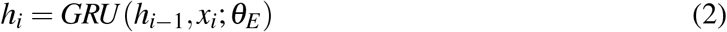

Two dense connection layers are then used to learn the mean vector *µ*_*z*_ and the standard deviation vector *σ*_*z*_. latent variable z is then sampled from the Gaussian distribution 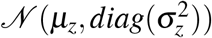

### Variational Attention

Attention mechanism tries to dynamically align 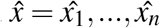 during generation. During decoding, the attention mechanism for step j of the decoder is computed between all the hidden states of encoder and hidden state at step j of decoder as :

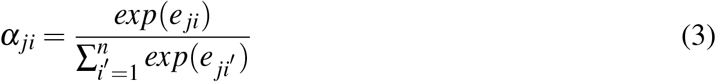

In the above equation *e*_*ji*_ is the pre-normalized score calculated as 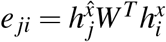 where 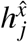 and 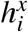 are the *j*’th and *i*’th hidden representation of decoder and encoder hidden states and W is a bilinear term to capture specific relation. The attention vector is then calculated by a weighted sum as:

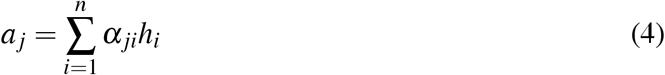

The posterior 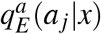 is modeled as another gaussian distribution 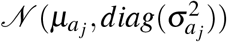 which is written as:

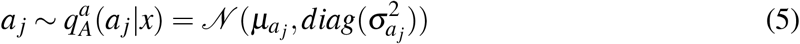

### Decoder

The decoder is a single layer GRU. The decoder at each step is provided with the latent space of encoder, attention vector computed at each step and the true input sequence 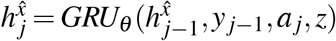

A softmax function at the end is used to predict the next word in the sequence 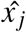 given the hidden representation at 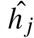:

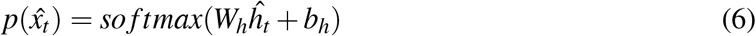

The loss function of the model with attention at each step of decoder can be written as:

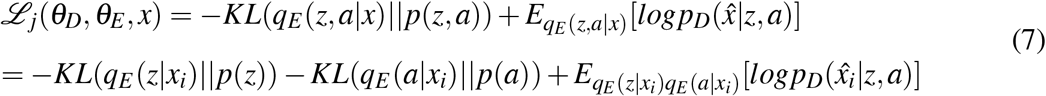

In the above equations *KL* is the Kullback-Leibler divergence between two distributions. The posterior *q*_*E*_ (*z, a*|*x*) = *q*_*E*_ (*z*|*x*)*q*_*E*_ (*a*|*x*) is factorized into two distributions since *a* and *z* are conditionally independent given x. The sampling can then be performed separately for *a* and *z*. The overall objective of the VAE with variational attention can then be written as:

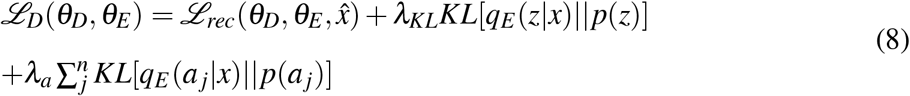

hyperparameters *λ*_*KL*_ and *λ*_*a*_ are the weights on the KL and attention terms of the loss function. Annealing is done on the *λ*_*KL*_ weight while the *λ*_*a*_ weight is kept constant. We used a monotonic annealing scheduler for training from 0 to maximum weight of the KL term.

## Results and discussion

### Antimicrobial prediction

To assess the quality of the generated peptides in the generative network, we first trained a binary classifier network for prediction of antimicrobial peptides. The training data for our model consists of AMPs and non-AMPs collected from multiple datasets as described in the methods section. The total dataset has 16,808 positive and 16,808 negative (non-antimicrobial) sequences. Our model architecture (figure 1A) consists of an Embedding layer, a convolutional layer, a bidirectional LSTM, a context attention and a sigmoid at the end to compute the probability of antimicrobial class for sequences. Attention mechanism is inspired by the the brain’s ability to prioritize segments of information during textual or visual processing. A bi-directional LSTM is a variant of RNN which encodes positional information from the sequence in a recurrent manner in both forward and backward directions. The context-vector attention generates a vector summary of all hidden states of the bidirectional LSTM using a weighting average, where the weights are learned during training. During training we used early-stopping to stop the training when there is no improvement after 5 consecutive training epoch. For training this model we performed a train/test split (70%/30%). For the classification model, we used a batch size of 32, a learning rate of 0.001, embedding dimension of 64, hidden dimension of 64. The number of convolutional kernels in the convolution layer were 64 with a size of 3. We also employed dropout with a dropout rate of 0.3 to avoid overfitting. The training and validation accuracy during training is shown in figure 1B. This model achieved an accuracy of 93.5% under a 10-fold cross validation. The accuracy of our model is comparable to other AMP prediction models such as AMPlify^30^ and ACEP^31^ which use deep learning models and report accuracies of 92.79% and 91.16% (for sequences less than 30) respectively.

### Training the generating network

For the generative model, we only used the AMP dataset consisting of 16,808 known AMP sequence. For training the generative VAE with variational attention, our architecture consisted of 128 hidden units for the Encoder and Decoder which are both single directional LSTM networks, a latent space dimension of 32. The model was trained for 100 epochs. The training process takes about 3 hours on a Tesla V100 GPU. During generation we experimented with different sampling methods such as Beam-search, Temperature sampling, Top-K sampling and Top-P sampling. Top-K sampling gave better results than other sampling methods with *K* = 5. The generative model architecture is shown in figure 2.

**Figure 2:**
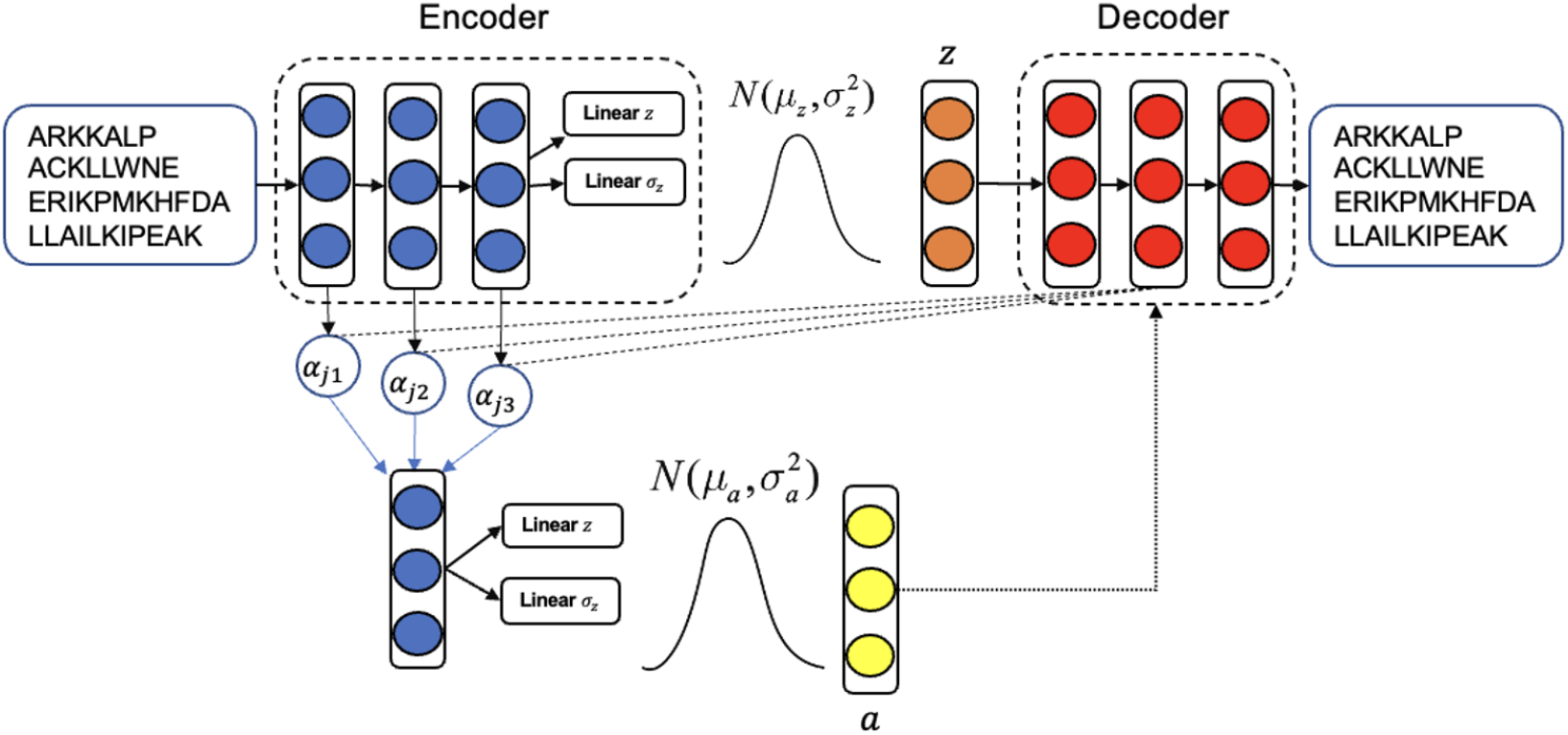
Illustration of AMP generative model with Encoder, Decoder and variational attention parts

During the training process we tokenize the sequences of amino acids into all the two natural amino acids and three additional tokens representing the start of the sequence “<SOS>“, the end of a sequence “<EOS>“, the padding “<PAD>“. In the word-dropout technique, we randomly replace an amino acid with “<UNK>” token during training. Since the peptides have different length, we added a padding token to the sequences so that they all have a fixed length of 30 during training and evaluation. The standard VAE where the weight on the KL term is 1, suffers from a KL vanishing problem which leads to i) an encoder that produces posteriors almost identical to Gaussian prior for all samples and ii) Decoder ignores the latent variable and the model reduces to a simple RNN Encoder-Decoder. Bowman et al.^19^ proposed two approaches to deal with this problem. A word-dropout which randomly replaces the words in sequences with an unknown token “<UNK>” during training to avoid overfitting in the model. And the second approach I KL-annealing where at the start of the training process, the weight of the KL term is small where z is learned to capture useful information for reconstructing x during training. Then the KL-weight increases monotonically to a maximum value. During training we used a monotonic annealing where the weight increases from a small value (0.001) to a maximum value. Not annealing the KL-term led to a posterior collapse and an uninformative latent space. We experimented with different weights on the KL and attention KL term and the results of generative model evaluation are shown in figure 3. For each set of hyperparameters we generated 10,000 peptide sequences from the generative model. The AMP-prediction model was used to predict what fraction of the generated AMPs are in fact antimicrobial. As shown in figure 3, increasing the *λ*_*KL*_ increases the accuracy of the generative model in generating antimicrobial peptides. In most *λ*_*KL*_ values, increasing *λ*_*a*_ also increases the accuracy of AMP prediction. BLEU is another metric used here to evaluate the generated peptides. This was originally proposed for evaluation of machine translation tasks by comparing the similarity between the translated sentences by the model and the true human references. Here we used BLEU score to compare the generated AMPs with the training data of real AMP sequences. BLEU score is calculated for every sample in the generated dataset *S*_*gen*_ against all the real AMP references *S*_*re f*_ as :

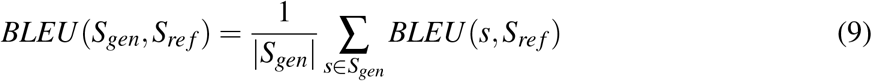

**Figure 3:**
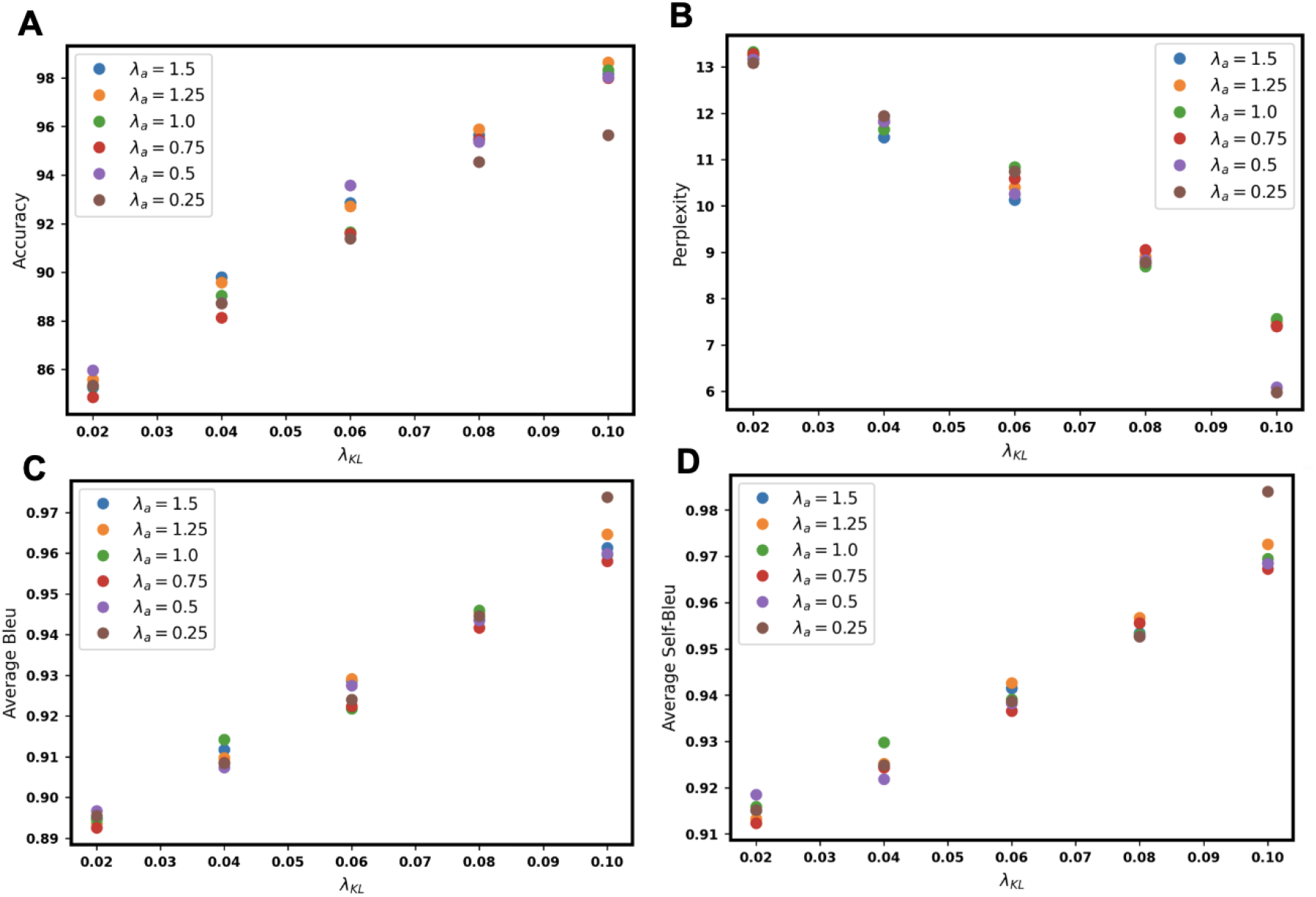
Evaluation of the AMP-generative model over different values of *λ*_*KL*_ and *λ*_*a*_ a) Accuracy of the generative model over 10,000 generated sequences using the trained AMP-prediction model B) average perplexity of the generated sequences using an external language model C) Average BLEU score (BLEU-2 to BLEU-5) of generated sequences D) Average Self-BLEU score (BLEU-2 to BLEU-5) of generated sequences

Higher BLEU score implies more overlap of n-grams in the generated data and the real AMPs. BLEU score for VAE without annealing, VAE with annealing and *λ*_*KL*_=0.08 and VAE-attn with different combinations of *λ*_*KL*_ and *λ*_*a*_ are calculated in table 1.

**Table 1.** BLEU, accuracy and perplexity of a few selected models

| Model | BLEU-3 | BLEU-4 | BLUE-5 | BLEU | ACC | PPL |
| --- | --- | --- | --- | --- | --- | --- |
| VAE (no anneal) | 0.999 | 0.984 | 0.911 | 0.9735 | 99.5 | 5.3 |
| VAE ($\lambda=0.08$) | 0.998 | 0.962 | 0.826 | 0.9465 | 96.2 | 8.09 |
| VAE-attn ($\lambda_{KL}=0.04$,<br>$\lambda_a=0.5$) | 0.997 | 0.921 | 0.711 | 0.90725 | 88.7 | 11.82 |
| VAE-attn ($\lambda_{KL}=0.04$,<br>$\lambda_a=1.5$) | 0.998 | 0.928 | 0.722 | 0.912 | 89.8 | 11.48 |
| VAE-attn ($\lambda_{KL}=0.06$,<br>$\lambda_a=0.5$) | 0.998 | 0.943 | 0.769 | 0.9275 | 93.6 | 10.26 |
| VAE-attn ($\lambda_{KL}=0.06$,<br>$\lambda_a=1.5$) | 0.998 | 0.944 | 0.773 | 0.92875 | 92.84 | 10.13 |
| VAE-attn ($\lambda_{KL}=0.08$,<br>$\lambda_a=0.5$) | **0.998** | **0.957** | **0.818** | **0.94325** | **95.6** | **8.8** |

### External language model

In NLP, given a sequence of words *x* = (*w*_1_, …, *w*_*n*_), language models estimate the probability distribution *P*(*x*) over it. A popular choice of language modeling is autoregressive model where:

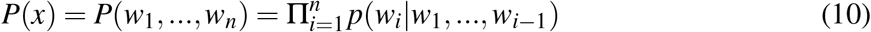

The likelihood of a sequence of words *P*(*x*) can be used as a proxy for its quality. RNNs and LSTMs are common architectures for autoregressive modelling. RNNs are trained to predict next word given the current word *w*_*i*_ and the hidden state of previous word *h*_*i*−1_ which is equivalent to maximizing the marginal likelihood *P*(*x*) of a sequence of words in the training data. RNN language models have been proposed to compute the perplexity of generated text which is a measure of fluency of machine generated text.^32^ we trained an LSTM language model on the AMP dataset. For this we split the training set into train (70%) and heldout or test (30%) set. Then we trained a character-level LSTM on the training set and calculated the perplexity on the heldout dataset. Our best model achieved a perplexity of 6.0 on the heldout dataset. Figure 3B shows the change in perplexity upon changing the *λ*_*KL*_ and *λ*_*a*_ values in VAE-attn model. Higher *λ*_*KL*_ gives a higher perplexity to the model. Table 1 shows the perplexity of a standard VAE (without annealing), a *β* -VAE with *λ* =0.08 and other selected models. A standard VAE has the lowest PPL which is even lower than that evaluated on the validation set. This shows that the standard VAE has collapsed to a denoising autoencoder which is just reconstructing back the original data.

In order to evaluate the diversity of the generated AMPs we used self-BLEU score which assesses the similarity between every generated sequence and the rest of the generated dataset. Lower self-BLEU score implied higher diversity of the generated text. The BLEU and self-BLEU scores for different *λ*_*KL*_ and *λ*_*a*_ weights are shown in figure 3C,D. Increasing *λ*_*KL*_ increases the BLEU and self-BLEU scores. We noticed that at higher *λ*_*KL*_ values the difference between different *λ*_*a*_ values are more noticeable. The comparison of self-BLEU for real AMPS, standard VAE (no annealing), *β* -VAE (*λ* =0.08) and other selected models is shown in table 2. The self-BLEU for real-AMPs is 0.968. This high value is due to the choosing a small cutoff (0.35) for removing repetitive AMPs in the data curation process which was chosen to maximize the number of selected AMPs for the generative model. Standard VAE without annealing shows a higher self-BLEU than the real AMPS which shows a very low diversity of generated sequences. *β* -VAE also shows a higher self-BLEU than other VAE-attn models. The KL divergence for each model was also calculated against the validation AMP dataset. A higher KL implies a higher difference between real and generated AMP distributions. However, a very high KL could lead to the model only generating random sequences. A comparison of KL for different models is shown in Table 2. Standard VAE have a KL of 0 which shows the posterior collapse and the model becoming a RNN language model. The KL for *β* VAE is 1.1 which is also lower than VAE-attn models which points to lower divergence more similarity of *β* VAE generated sequences to the real AMPs.

**Table 2.** self-BLEU (sBLEU) for 3,4 and 5-grams and KL divergence

| Model | sBLEU-3 | sBLEU-4 | sBLUE-5 | sBLEU | KL |
| --- | --- | --- | --- | --- | --- |
| **real AMPS** | 0.998 | 0.967 | 0.909 | 0.968 | - |
| **VAE (no anneal)** | 0.998 | 0.984 | 0.911 | 0.973 | 0 |
| **VAE (lamda=0.08)** | 0.998 | 0.962 | 0.826 | 0.946 | 1.1 |
| **VAE-attn ($\lambda_{KL}=0.04$,**<br>**$\lambda_a=0.5$)** | 0.999 | 0.929 | 0.764 | 0.923 | 13.1 |
| **VAE-attn ($\lambda_{KL}=0.04$,**<br>**$\lambda_a=1.5$)** | 0.994 | 0.9333 | 0.771 | 0.924 | 12.4 |
| **VAE-attn ($\lambda_{KL}=0.06$,**<br>**$\lambda_a=0.5$)** | 0.993 | 0.941 | 0.819 | 0.938 | 7.0 |
| **VAE-attn ($\lambda_{KL}=0.06$,**<br>**$\lambda_a=1.5$)** | 0.994 | 0.944 | 0.828 | 0.942 | 6.4 |
| **VAE-attn ($\lambda_{KL}=0.08$,**<br>**$\lambda_a=0.5$)** | **0.998** | **0.957** | **0.818** | **0.943** | **3.2** |

We also investigated the generated antimicrobial peptides through their physicochemical properties such as length, charge, hydrophobicity and hydrophobic moment in figure 4. Specifically the generated sequences are rich in amino acids such as Lys, Leu, Arg, Ala, Ile, Gly, Val, Phe and Trp in that order. As shown in the distribution of charges for generated peptides, most of them have a positive net charge due to presence of Lys and Arg residues. Furthermore, the hydrophobic moment shows that most of these are *α*-helix peptides. These observations highlight the close properties of the generated peptides with the dataset of real antimicrobial peptides which point to the fact that the generated peptides have antimicrobial activities.

**Figure 4:**
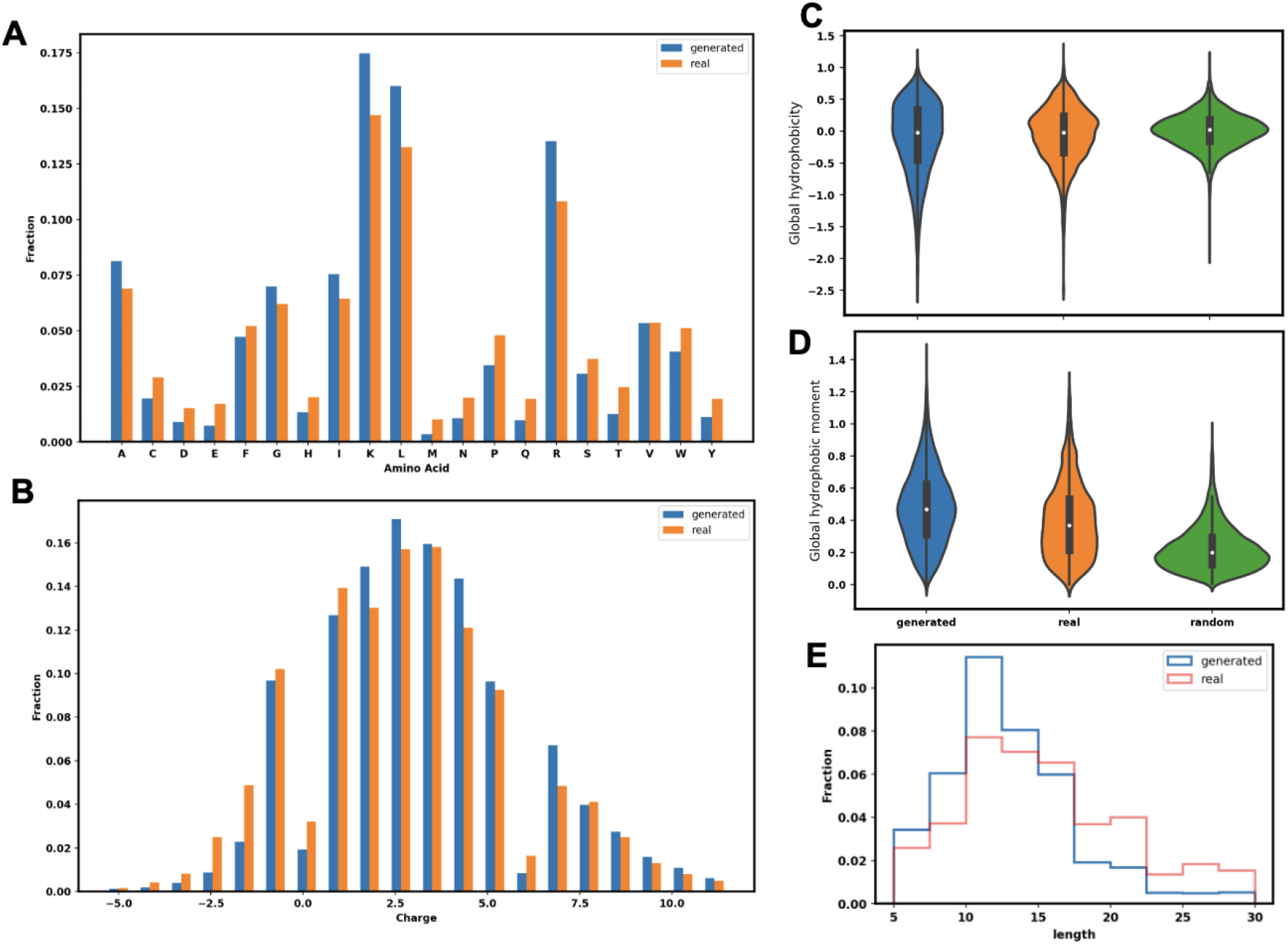
Physico-Chemical properties of the generated 10,000 AMP sequences and their comparison with real AMP dataset. A)fraction of different type of amino acids B) distribution of charge in real and generated AMPs C) Global hydrophobicity over generated, real and random sequences Global hydrophobic moment for generated, real and random sequences E) Sequence length distribution for generated and real AMPs

## Conclusion

Antimicrobial peptides have shown great potential as alternative therapeutics for bacterial resistance. In this study we use deep learning generative model attention based variational autoencoder to generate novel and high quality sequences of AMPs. A bypassing phenomena has been observed when using deterministic attention in a VAE framework. The bypassing phenomena makes the latent space uninformative and sampling from this space would give random results during generation. Since attention has been shown to improve tasks in NLP, it is tempting to include attention mechanism in our model. But a deterministic attention is not possible for sequence generation task on the same dataset so we opted for a variational attention mechanism. Therefore, Bahuleyan et al.^23^ proposed a variational attention approach where the attention vector is modeled as random variable by imposing a prior Gaussian. Here we used variational attention based variational autoencoder to generate novel AMPs. The generated AMPs from our best model were evaluated using an antimicrobial prediction network which showed a more than 95% probability. We have also used evaluations metrics such as BLEU, self-BLEU and perplexity which showed that high quality of the generated sequences. Moreover, we compared the physicochemical properties of the generated peptides with the real AMPs which showed closeness of these properties. The future direction of this work will be using post-evaluation models such as regression models to predict antimicrobial (MIC) activity of the generated peptides and a toxicity prediction to further choose a few selected peptides. Further evaluation of the generated peptides can be performed by performing MD simulations and experimental validation.

## Acknowledgement

This work was partially supported by the National Heart, Lung and Blood institute at the National Institute of Health for B.R.B. and M.G.; in addition, it was partially supported by the National Science Foundation (grant number CHE-2029900) to J.B.K. The authors acknowledge the biowulf high-performance computation center at National Institutes of Health and Lobos for providing the computational time and resources for this project.

## Notes

### Competing Interest Statement

The authors have declared no competing interest.

